# Transcription promotes discrete long-range chromatin loops besides organizing cohesin-mediated DNA folding

**DOI:** 10.1101/2023.12.29.573667

**Authors:** Christophe Chapard, Nathalie Bastié, Axel Cournac, Olivier Gadal, Romain Koszul, Frédéric Beckouët

**Affiliations:** Institut Pasteur, CNRS UMR 3525, Université Paris Cité, Unité Régulation Spatiale des Génomes, 75015 Paris, France; Molecular, Cellular and Developmental biology department (MCD), Centre de Biologie Intégrative (CBI), Université de Toulouse, CNRS, UPS, 31062, Toulouse, France

## Abstract

The multi-layered arrangement of eukaryotic genomes and chromosome spatial organization dynamics are of functional importance for gene expression, DNA replication and segregation. SMC complexes are essential instruments of chromosome folding by carrying out long range intra-chromatid DNA looping. Cohesin, in addition to tether sister chromatids, also ensures dynamic regulation of gene expression in mammals by promoting interaction between distal regulatory elements and promoters whereas transcription affects genome folding in numerous organisms and in multiple ways. Here, we comprehensively dissect the relative contributions of transcription and cohesin complexes, as well as their interplay, on the yeast *S. cerevisiae* genome organization through DNA borders and loops. Transcription activation specifically induces appearance of DNA borders and loops, independently of SMC complexes, while also directly interfering in addition with cohesin-mediated loop expansion.

## Introduction

Chromosomes exhibit a three-dimensional (3D) organization that can affect or regulate DNA processes including gene regulation, DNA repair, or segregation (1–3). This organization consists in an intertwined network of structures such as long-range DNA loops, large self-interacting chromosomal domains up to hundreds of kilo-bases, or also compartmentalization of chromosomal regions (4–6). These layers of organization are influenced, or regulated, by molecular processes involving for instance topoisomerases, transcription or the ubiquitous structural maintenance of chromosome (SMC) complexes (cohesin, condensin, Smc5/6) (7–12). Understanding the relative contributions of each factor to establish, maintain, and/or regulate chromosome functional organization remains challenging (13).

The loop extrusion (LE) model proposes that SMC rings organize genomes by gradually enlarging small DNA loops into larger structures until they encounter a roadblock and/or a release signal (4,14–17). CCCTC-Binding Factor (CTCF) were identified as major roadblocks to LE in mammals (3,11,18) while other roadblocks have also been described, especially in organisms lacking CTCF, such as centromeres, telomeres or sister chromatid cohesion in yeast (19,20)(21).

The findings that cohesin localizes between genes that are transcribed in a convergent direction in organisms as different as *Saccharomyces. cerevisiae*, *Schizosaccharomyces pombe* and in mammals depleted for both for CTCF and Wpl1, suggest that active transcription may also displace or stop DNA loop extruding cohesin, and/or interfere with cohesin loading process (22–26). Recent studies suggest that transcription negatively regulates cohesin dependent DNA loops (12,27–30). CTCF/Cohesin independent Enhancer-Promoter contacts, that may require PolII for their formation, have also been recently described in mammals (10,12,28,31). Overall, these studies highlight the interplay between active transcription and cohesin-mediated LE in eukaryote functional genome organization, and probably more generally in all kingdoms of life given the ubiquity of these molecular processes and complexes (32). However, how these processes articulate with each other remains unclear. It is therefore essential to improve our understanding of the regulations at play in order to fully comprehend chromosome functional organization.

In this work we precisely dissect the causal relationships between these partners, and disambiguate large aspects of the regulation of chromosome folding in yeast, using cell-cycle synchronization, inducible transcription, and controlled depletion of members of SMC complexes that allow monitoring of the progressing DNA loop expansion. In particular, we identify the SMC-independent existence of transcription-dependent *de novo* DNA loops, bridging for instance a highly expressed gene to active neighbouring genes *in cis*. Furthermore, transcription activation also results in the formation of strong DNA borders that prevent or restrict the establishment of cohesin-dependent DNA structures, such as metaphase chromatin loops, while having little impact on the maintenance of these structures once formed. Finally, we demonstrate that transcription-dependent DNA borders can also act as semi-permissive barriers to dynamic cohesin mediated DNA loop expansion. Altogether, our results reveal some important and respective contributions of two of the major, ubiquitous players of chromosome folding, transcription and the SMC cohesin.

## Results

To explore the effect of transcription induction on chromosome 3D organization, we looked at the effect of galactose-inducible activation of the well-studied *GAL7-GAL10/GAL1* locus (*GAL7-10/1*) of *S. cerevisiae* on chromosome II (**Fig. 1a**, **Methods)**. The genome organization and RNA polymerase II profiles of cells grown either in presence of glucose or galactose and synchronized at the G1 stage of the cell cycle were characterized using Hi-C and ChIP-seq, respectively (**Methods**) (22). As expected, upon growth in galactose, the locus was highly enriched in PolII compared to cells grown in glucose (**Fig. 1b**). Both the ratio of induced vs. uninduced maps (**Fig. S1a**) and comparison of the normalized maps (**Fig. S1b**) show that several changes accompany the gene induction. Firstly, the activated locus appears as a strong boundary to cis-DNA contacts that otherwise join either side of the inactive locus together (**Fig. 1b, S1a**), resulting in a visual partitioning of the chromosome into sub-domains. Other GAL loci of the genome, which are also induced in presence of galactose, display similar patterns (**Fig. S1b**). Along the DNA border, dotted contact pattern also appears at the level of the induced *GAL7-10/1* genes, pointing at the formation of chromatin loops that bridge the locus with neighbouring regions (**Fig. 1b**). These loops differ from G2/M cohesin-mediated loops in at least three ways: 1) cohesin deposition along chromosomes is reduced in G1 (33), 2) they persist along G2/M metaphase chromosomes depleted for Scc1 (**Fig. S2f**), and 3) their bases do not overlap with convergent genes (19,24) (**Fig. 1c lower panel**). A virtual 4C analysis also shows enrichment in contacts between the induced GAL locus and actively transcribed, PolII-enriched regions up to one hundred kb away (**Fig. 1c upper panel, S1c**). These new loops associated with PolII enrichment can cross the centromere to bridge regions on both chromosome arms, another stark difference with G2/M cohesin-mediated DNA loops for which the centromere represents a seemingly un-crossable boundary (19).

**Figure 1:**
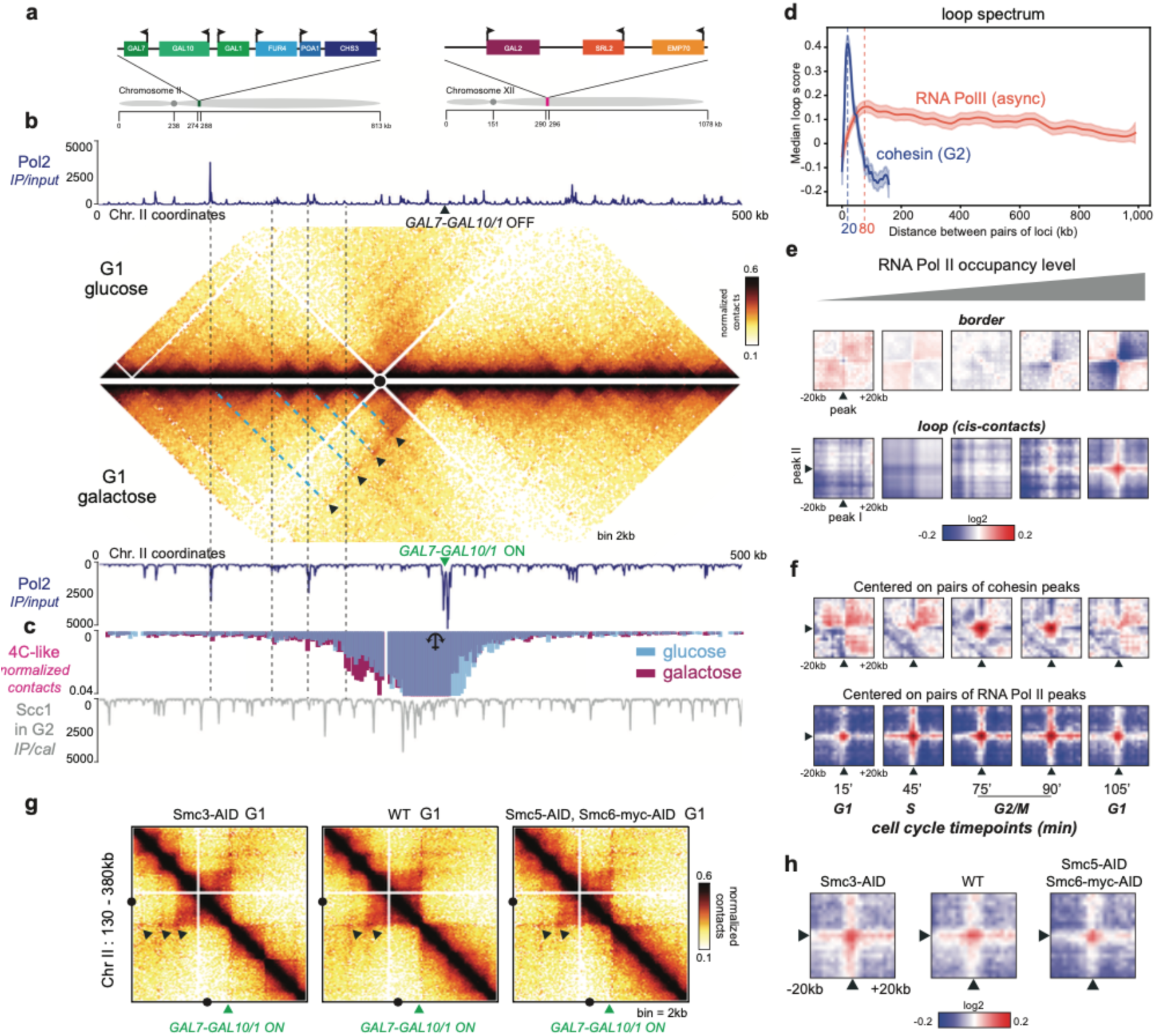
Active Pol2 delimits chromatin domains and induces discrete long-range DNA loops. a) Schematic representation of *GAL* loci. b) Hi-C contact maps (bin: 2kb), and corresponding RNA Polymerase II ChIP-seq profiles, of a section of chromosome II containing the GAL7-10/1 locus in cells synchronized in G1 in either glucose (top) of galactose (bottom). Loops with the GAL7-10/1 locus as one basis are indicated with black triangles. For each loop, the coordinate of the other basis is identified following the blue, then dark, dotted lines. c) Upper panel: A virtual 4C plot with a viewpoint (anchor) just upstream the GAL7-10/1 locus of cells in either glucose (pink) or galactose (light blue) is shown below RNA polymerase II Chip-seq profile in galactose. Lower panel: the Virtual 4C plot a cohesin (Scc1) ChIP-seq profile of G2 arrested cells is shown in grey. d) Loop spectrum showing the loop score computed by Chromosight as a function of the genomic distance separating 2 Pol II RNA peaks . e) Pileups of RNA Pol II peaks at the positions sorted by RNA pol II levels (5 groups) along the diagonal (top) and at long distances (60 kb – 460 kb) (bottom). f) Pile up plot for cohesin loops (pairs of Scc1 peaks separed by genomic distances between 10 kb and 50 kb) and for RNA Pol II loops (pairs of RNA Pol II peaks at long distances (60 kb – 460 kb)) at different time points during the mitotic cell cycle. g) Hi-C contact maps (bin: 2kb) of the chromosome II region containing the GAL7-10/1 locus following depletion of SMCs by IAA addition. h) Pile up plot for RNA Pol II enriched regions over long distances (60 kb – 460 kb) for WT and SMC depletion (same data as panel g).

We then explored whether contacts between active genes could be detected genome wide. To do so, we applied the “*quantify*” mode of the program Chromosight to cells arrested in G2/M in medium containing glucose. This mode computes “loop scores” reflecting an enrichment/depletion in contacts between pairs of loci, and plots the corresponding loop spectrum as a function of the genomic distance between them (see **Methods**)(34). As a reference, we also computed the *quantify* loop spectrum using pairs of loci in cis enriched in cohesins in G2 (**Fig. 1d**, blue curve) (19). The spectrum displays the expected maximum for genomic distances around 20 kb, corresponding to the average loop sizes at this stage. When applied on pairs of loci enriched in RNA Pol II in cis, the analysis reveals an enrichment in contacts over very long distances, up to 1 Mb, with a maximum score around 80 kb (**Fig. 1d**, red curve), showing that active genes have a slight tendency to contact each other in cis more than other regions. To determine further the correlation between the levels of gene activation and the apparition of transcription-dependent DNA borders and loops, we computed pile-up contacts maps of 40 kb windows centred on RNA Pol II peaks classified according to their Pol II occupancy levels (**Methods**; **Fig. 1e**). Both patterns appear stronger as the sites are strongly enriched in Pol II, in agreement with previous observations from bacteria or humans (34–36). Finally, enrichment of contacts between long-distance RNA pol II peaks is detectable throughout the cell cycle, in contrast to cohesin dependent DNA loops which only form only during S and G2/M phases (**Fig. 1f**).

Altogether, these results show that upon transcription activation creates a contact boundary in the map sometimes associated with punctate contacts to neighbouring actively-transcribed regions.

### Transcription mediated DNA structures are independent of SMC complexes

Members of the SMC family are involved in the formation of long-range chromatin contacts. We therefore explored whether transcription-mediated chromatin structures depended in one or the other SMC complexes present during G1, namely cohesin and SMC5/6; but not condensin that is inactivated through the natural degradation of the essential Ycg1 subunit up to undetectable levels (37). We therefore performed controlled degradation of either Smc3-aid (cohesin) or Smc5-aid/Smc6-aid in cells that also express the *GAL4-ER-VP16* fusion protein, allowing inducible activation of GAL promoters in response to oestradiol, even in the presence of glucose (38) (method) (**Fig. S2a, S2b, S2c et S2d**). The direct comparison of cells grown in identical culture conditions is therefore granted, as opposed to glucose *versus* galactose treated cells (**Fig. S2a, S2b, S2c et S2d**). Hi-C contact maps were generated from cells arrested in G1, depleted from either cohesin or Smc5/6, and with GAL genes induced by oestradiol (**Fig. 1g, S1d, S1e and S1f; Methods**). Hi-C maps showing transcription induced DNA border appearance at the GAL locus in all conditions and in a manner that was barely undistinguishable from WT control (**Fig. 1g**). We further noticed that the signal was even neater after removal of cohesin, leading to the appearance of other DNA borders at different loci (**Fig. 1g, Fig. S2e**), suggesting that residual cohesin binding to DNA in WT cells in G1 could inhibit either transcription activation, DNA border formation, or both. Finally, we computed the genome wide pile-up in the cells in these cells showing that the signal is of the same intensity in absence of these two SMCs (**Fig. 1h**).

Overall, active transcription mediates DNA borders and long-range loops in G1 through a mechanism that is independent of individual SMC complexes. This observation confirms that transcription per se has prominent effects on chromosome 3D organization.

### Transcription restricts the formation of cohesin-dependent intrachromosomal contacts

We next sought to decipher the nature and causative relationship(s) between transcription and the formation of the chromatin loops mediated by cohesin that decorate yeast chromosomes during G2/M (19,39). To look at the effects on 3D organization, WT cells growing exponentially in glucose medium were arrested in G1 using alpha-factor, and GAL genes induced using oestradiol (**Methods**). Cells were released from G1, and allowed to progress into S-phase **before** being arrested in G2/M with nocodazole (**Fig. 2a and 2b**). Hi-C contact maps, and Scc1 and RNA PolII ChIP-seq, were generated for the different time points in presence or absence of oestradiol induction of GAL genes. The normalized contact maps and average contact frequencies as a function of genomic distance curves P(s) show that oestradiol does not disturb the overall establishment of the overall genome organisation (**Fig. 2c – e and Fig. S2c-d**). However, local changes occurred at the level of galactose inducible genes *GAL2* and *GAL7-10/1* (**Fig. 2d, e**). Previous work has shown that induced transcription at the chr. XII *GAL2* locus in galactose-grown cells prevented the cohesin accumulation observed on the gene body in G2/M when cells are grown in glucose (24) (loci in **Fig. 1a**). Instead, cohesin enrichment was observed at a downstream place, at the 3’ end of the active *GAL2* gene (22). The effect of this repositioning of cohesin on the 3D organization of chromatin had remained unknown until now. Our new data reveal that cohesin repositioning at this locus also corresponds to the repositioning of the base of the associated DNA loop (**Fig. 2d, f**) leading to a decrease in loop size by about 2kb. This result demonstrates that transcription restricts cohesin-dependent loop positioning.

**Figure 2:**
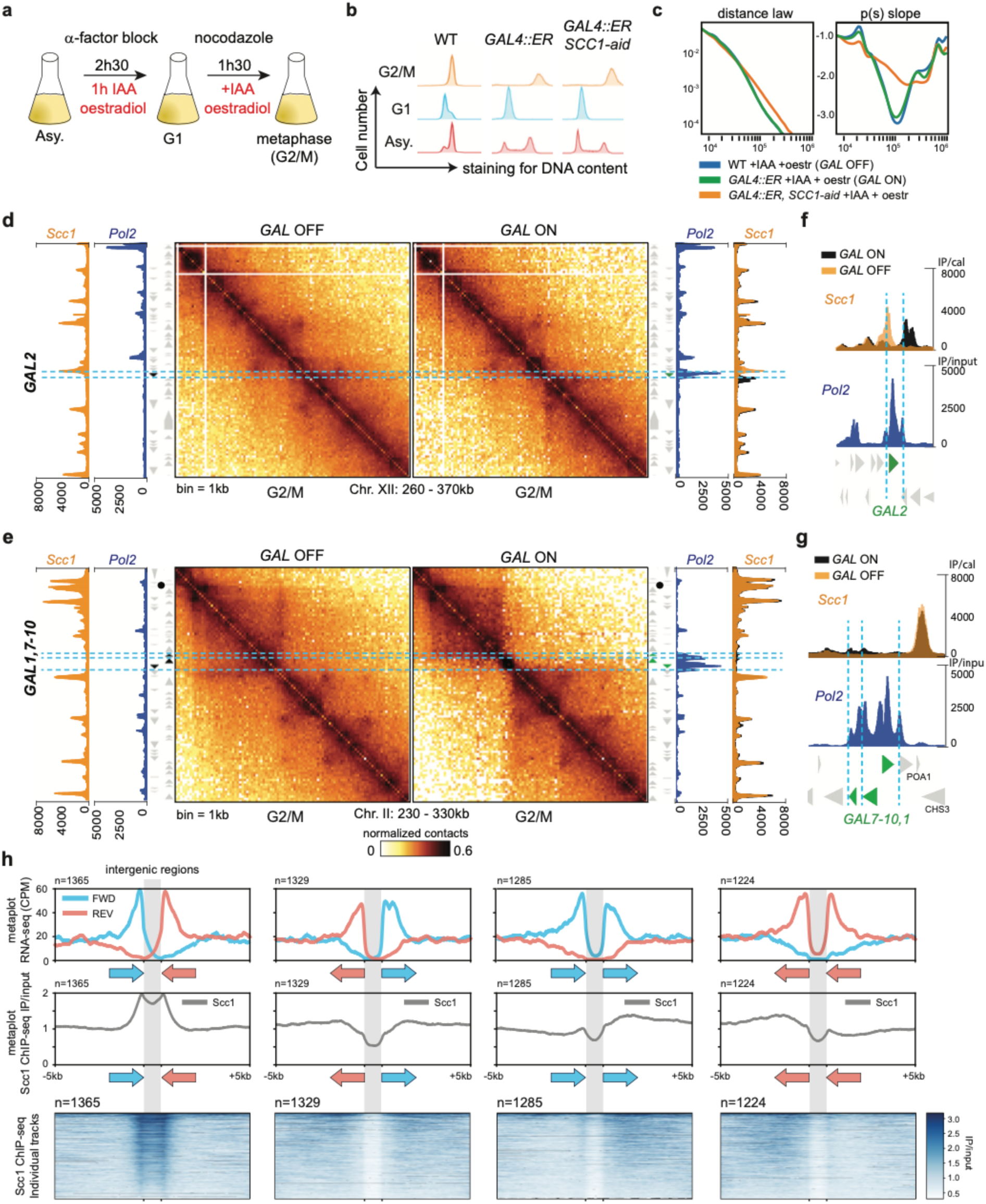
Transcription dependent DNA borders hinder establishment of cohesin mediated DNA structures. a) Schematic representation of the experimental protocol used to process yeast cells from G1 to metaphase, concomitantly with oestradiol-induced GAL gene induction and auxin-mediated SCC1-AID depletion. b) Synchronisation is controlled with by flow cytometry. Asy: asynchronous. c) Contact frequency curves P(s) representing the average contact frequency as a function of genomic distance (bp), and their respective derivative curves. d) Hi-C contact maps (bin: 1kb) of a region of chr. XII containing the *GAL2* locus, with the corresponding RNA Pol 2 and Scc1 ChIP-seq profiles. Cells are synchronized in metaphase, without (left) or with (right) oestradiol-induced activation of *GAL2*. e) Same as d) but for a region of chr. II containing the *GAL7-10/1* locus. f) Top: magnification of the overlaid Scc1 ChIP-seq profiles before (light orange) and after (black) oestradiol-mediated induction of *GAL2* on chr. II. Bottom: corresponding RNA Pol 2 ChIP-seq profile (blue). g) same as f) but for the region of chr. XII containing the *GAL7-10/1* locus. h) Average profiles of Scc1 ChIP-seq signal measured (33) across intergenic regions ±5 kb classified into 4 groups according to the orientations of the flanking genes (forward and reverse) .

In contrast to inactive *GAL2*, the silent *GAL7-10/1* is devoid of cohesin enrichment. Nevertheless, a stripe emerged 3.8 kb away from the inactive *GAL7-10/1* region at the closest convergent genes (*POA1* and *CHS3*), and extends up to the centromere, 38 kb upstream (**Fig. 2e**; Dotted yellow line). This stripe, that reflects enriched contact in-between this position and all positions up to the centromere, disappears when Scc1 is depleted and is therefore cohesin dependent (**Fig. S2f**). The *GAL7-10/1* locus lies in-between the anchor points that form this cohesin dependent folding. Oestradiol activation of *GAL7-10/1* prior to replication and G2/M arrest strongly modified the contact pattern (**Fig. 2e, Fig. S2c)**. Indeed, the induction of the 6,2 kb long track, confirmed by the large RNA PolII enrichment (**Fig. 2g**), abrogates apparition of the stripe pattern.

In addition, calibrated ChIP-seq analysis revealed slight but significant cohesin enrichments at the 3’ end of the actively transcribed *GAL7*, *GAL10* and *GAL1* genes, compared to inactive conditions (**Fig. 2g**). In contrast to most of cohesin peaks that appear at sites of convergent transcription (19,24), the enrichments affected co-oriented genes (*GAL1*) or divergent (*GAL7-10*) genes. This result reveals that active transcription positions cohesin at 3’ end independently of the orientation of the neighbouring genes.

A genome-wide analysis of cohesin enrichment over pairs of genes classified according to their relative chromosomal orientation highlighted that this is a general phenomenon, with a clear tendency for cohesin to accumulate in 3’ positions of colinearly oriented genes (**Fig. 2h**). Altogether, our experiment suggests that transcribing polymerases have an active role on the regulation of cohesin dependent DNA folding, and appear not only to redistribute cohesin locally but also affect long range chromatin structuration by abrogating cohesin-mediated structure apparition (**Model, Fig. S2g**).

### Transcription is an active and semi-permissive barrier for loop expansion

Former works revealed that inactivation of Pds5 engages centromeric regions in long-range cis contacts with discrete loci along chromosome arms, leading to the establishment of very long DNA loops (CEN loops). CEN loops can stem from the chromosome arms and extend to the centromeres, and/or originate from centromeres and extend to the chromosome arms. To test whether transcription stops or slows the translocation process, we analyzed how an active *GAL7-10/1* locus directly interferes with *CEN* loop formation following the controlled depletion of Pds5. Pds5-AID cells expressing or not the *GAL4-ER-VP16* construct were arrested in G1 using alpha factor. Oestradiol was then added to both cultures and the cells released into S-phase, then arrested again in G2/M with nocodazole (**Fig. 3a, b**). Auxin was added to the medium to induce degradation of Pds5-AID (**Fig. 3a, c**). Following auxin addition, cells were sampled every 30 minutes for 2 h to monitor chromosome 3D organisation with Hi-C (**Fig. 3d and Fig. S3b**). Pds5-AID cells not expressing the *GAL4-ER-VP16* construct were used as controls. Pds5-AID depletion was achieved similarly in both conditions throughout the time course as assessed by the decrease of Smc3 acetylation (**Fig 3c**). At the same time, average contact frequencies as a function of genomic distance curves P(s) showed a progressive increase in long range DNA contacts (**Fig. S3b**). Normalized Hi-C maps, as well as 4C-like analysis using CEN2 as a genomic viewpoint, confirmed that Pds5 degradation in cells with silent *GAL7-10/1* engages *CEN2* in long-range, intra-arm contacts, resulting in CEN loops of increasing size (**Fig. 3d, e and Fig. S3a**). We verified using calibrated RNA PolII ChIP-seq that transcription was not affected upon Pds5-AID depletion. On the other hand, the pattern of *CEN2* contacts in the strain with *GAL7-10/1* activated prior to Pds5 degradation was very different. First, following Pds5 depletion DNA loops bridging *CEN2* with loci across *GAL7-10/1* were strongly attenuated (**Fig. 3e**). Large DNA loops increasing in size nevertheless gradually appeared along the arm, with the *GAL7-10/1* locus at their basis instead of CEN2 (**Fig. 3f**). This observation suggests that expansion of CEN loops is halted by the activated *GAL7-10/1* locus. This transcription-induced DNA loop roadblock to loop extrusion is semi-permissive as loops between *CEN2* and distant loci, bridged across the *GAL7-10/1* locus, gradually accumulate over time, albeit to later time points than on other chromosomal arms (**Fig. 3e**). Furthermore, the contacts made by *GAL7-10/1* engaged in DNA loops emanate from the same loci than that of regular CEN loops (**Fig. 3e, f**). Here too a closer examination of cohesin deposition profiles showed that Pds5-AID degradation promotes their enrichment at the 3’ end of *GAL7*, *GAL10* and *GAL1*, further supporting that active transcription tracks represent transient roadblocks to incoming cohesin, independently of the orientation of neighbouring gene (**Fig. 3g**).

Altogether, these data revealed that transcription activation represents a semi-permissive barrier for loop expansion (**Model Fig. S3c**).

**Figure 3:**
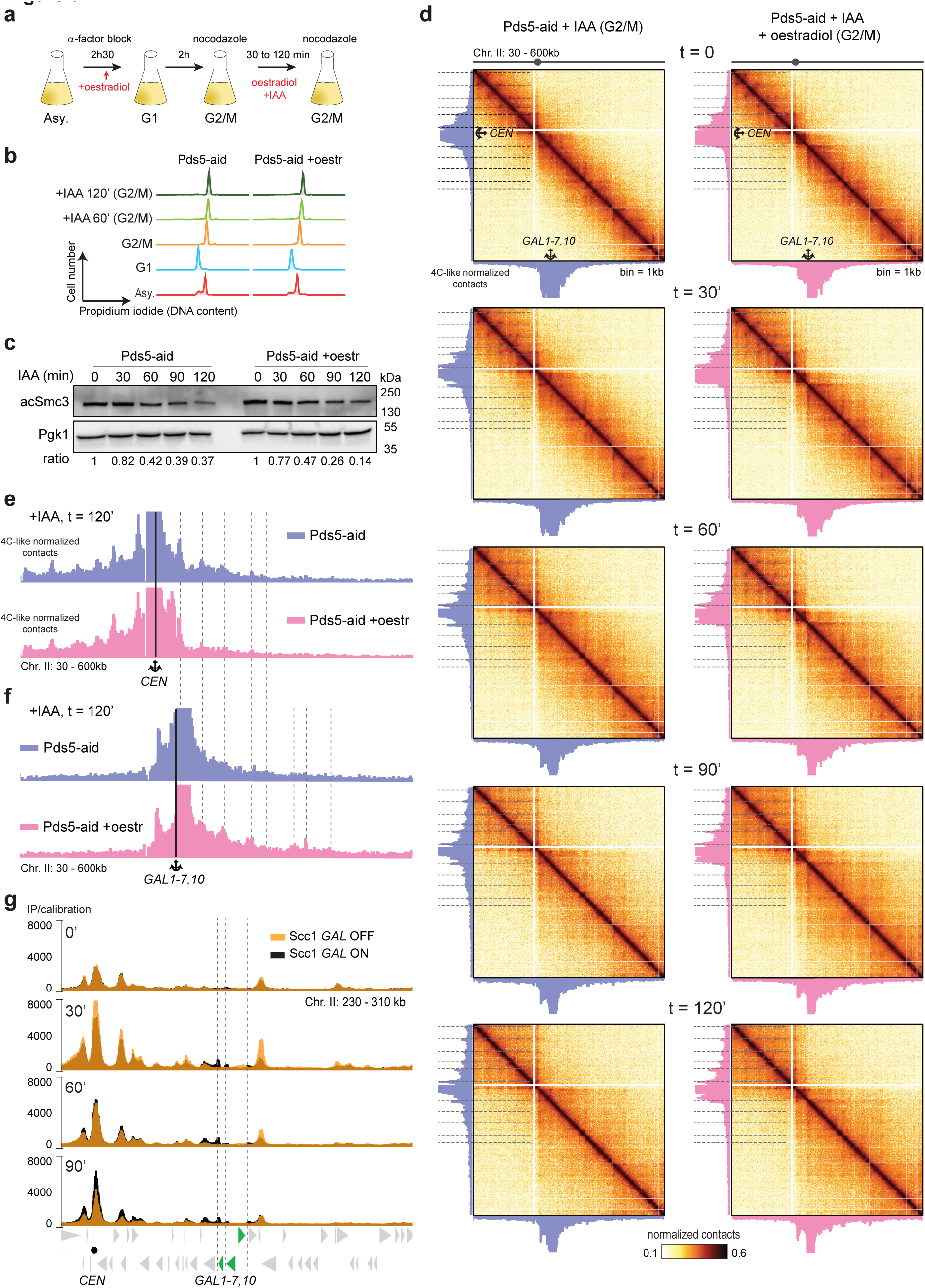
Transcription dependent DNA borders are semi-permissive barriers to cohesin mediated DNA loop expansion. a) Schematic representation of the experimental protocol for Pds5 auxine-mediated degradation in G2 in presence or absence oestradiol-induced GAL induction. b) Synchronisation is controlled with by flow cytometry until 120 minutes after the addition of IAA. Asy: asynchronous. G1: α-factor arrest; Meta.: metaphase; oe: oestradiol. c) Pds5 degradation was assessed by western blot through the Pds5-dependent Smc3 acetylation. Degradation was quantified relatively to the loading control Pgk1 (ratio). d) Hi-C contact maps (bin: 2kb) of a region (30-600kb) of chr. II during auxine-mediated Pds5 depletion (+IAA). The maps are ordered from top to bottom according to the time in presence of IAA. Time 0: no IAA, and then 30-, 60-, 90- and 120-min following IAA addition. For each map, two 4C-like profiles using either CEN (y axis) or *GAL7-GAL10/1* (x axis) loci as anchors are plotted. e) Magnifications of the 4C-like profiles using CEN as an anchor along chr. II in absence (Top blue) or in presence (Bottom pink) of GAL genes induction, 120 min after IAA-mediated Pds5 depletion. f) Magnifications of the 4C-like profiles using *GAL7-GAL10/1* as an anchor along chr. II in absence (Top blue) or in presence (Bottom pink) of GAL genes induction, 120 min after IAA- mediated Pds5 depletion. g) Overlaid Scc1 ChIP-seq profiles of a region (230-310 kb) of chr. II with (black) or without (yellow) GAL activation, and throughout Pds5 depletion (from top to bottom). CEN: centromere. Green arrow: *GAL7-GAL10/1* locus.

### Transcription interferes with, but does not prevent, the maintenance of cohesin dependent DNA structures

To analyse whether transcriptional activation can also interfere with the maintenance of wild type chromatin loops, we activated transcription **after** the establishment of cohesin-mediated loops during S phase. Cells were arrested in G2/M, the *GAL7-10/1* locus was induced by oestradiol, and cells processed with Hi-C (**Fig. 4a-d**). Hi-C maps revealed that transcriptional activation of *GAL7-10/1* while the cells are in G2/M results in a local DNA border, but with little if no impact on the maintenance of local cohesin-mediated structures that remain in place following induction (**Fig. 4d**). In addition, cohesin deposition at the activated locus remained also unchanged (**Fig. 4d**). Therefore, these experiments show that, while active transcription interferes with cohesin mediated DNA loops and structures during their formation, they are, once established in G2/M, largely unaffected by subsequent transcription activation. This observation argues against a strong cohesin *de novo* loading and/or highly dynamic translocation, at least locally at these active genes in G2/M of WT cells. Indeed, if the cohesin holding the bases of wild-type DNA loops were dynamic, one would expect that transcription activation would inhibit the *de novo* proper establishment of DNA loops and structures. This result therefore strongly supports our previous study indicating that a subpopulation of cohesin dependent loop is stable in mitosis (40).

**Figure 4:**
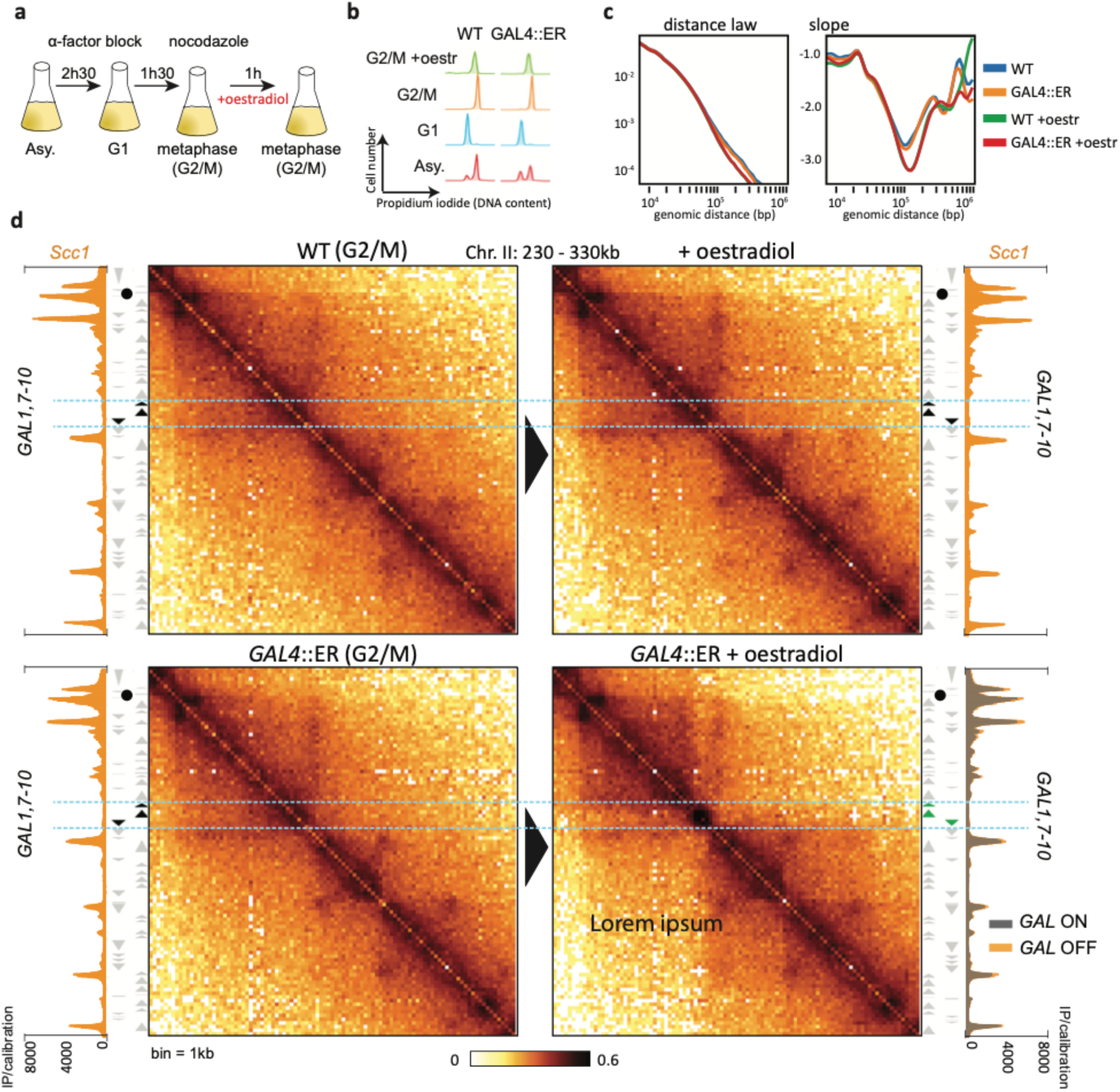
Cohesin mediated DNA structures are maintained upon appearance of transcription dependent DNA borders. a) Schematic representation of the experimental protocol followed for oestradiol-induced GAL induction during G2. b) Cells synchronisation was monitored by flow cytometry. Meta: metaphase; Asy: asynchronous; oe: oestradiol. c) Contact frequency curves P(s), representing the average contact frequency as a function of genomic distance (bp), and their respective derivative curves. d) Hi-C contact maps (bin: 1: kb) and Scc1 ChIP-seq profiles of a region (230-330kb) of chr. II showing the effect of the activation of the *GAL7-GAL10/1* locus during G2 on both chromatin loops and cohesins enrichment sites. Top: control WT cells. Bottom: GAL4-ER-VP16 expressing cells allowing the oestradiol-induced activation of GAL genes.

On the contrary, cohesin deposition at *GAL2* locus was strongly affected upon activation of transcription in G2/M, as previously described (22)(**Fig. S4a**). This change was associated with a widening of the associated DNA loop signal as seen on the Hi-C map, possibly due to partial repositioning of the loop anchor in a fraction of the cell population (**Fig. S4b**).

Thus, unless activation of transcription occurs immediately on the base of a DNA loop, cohesin dependent DNA structures and loops are maintained and resistant to the stress induced by transcription generated DNA loop borders in G2/M (**Model Fig. S4c**).

## Discussion

### Direct effect of transcription on chromatin long- and short-range folding

Both transcription and cohesin participate in genome folding, which in turn regulates DNA-related processes, including fine-tuned modulation of transcription itself (41,42). Here we disambiguate this interplay by revealing the respective contributions of transcription and cohesin to chromatin organisation. First, we show that induced GAL promoters lead to both DNA borders delimiting distinct domains and enriched contacts with neighbouring transcribed loci, resulting in long range DNA loops independently of individual SMC complexes. They may either result from direct clustering of RNA Polymerase complexes or transcription factors, or from indirect contacts resulting from redirection of the active regions to the same nuclear region, for instance in the nuclear periphery (43–45). Upon cohesin depletion, transcription mediated DNA folding appears stronger, suggesting cohesin folding suppresses, or at least masks in Hi-C map, these structures. Active transcription of native or artificial promoters also results in strong enrichment in local contacts along the transcribed tracks, resulting in boundaries in Hi-C contact maps whose precise nature remains unclear (46) but that are strongly reminiscent of transcription-induced domains observed along bacteria’s chromosomes (35) .

### Impact of transcription on cohesin-mediated chromatin loops

Several studies aimed at understanding how transcription influences formation of cohesin-dependent chromatin loops (12,27–30). In the present study, we precisely dissect the effect of GAL genes activation on cohesin mediated DNA loop formation, expansion and maintenance. First, we show that gene activation prevents or constraints formation of cohesin dependent DNA structures such as chromatin loops. At the Gal2 locus, DNA loops are restricted in size suggesting in addition that the basis of a DNA loop is defined by activities taking place on the DNA rather than specific DNA motives (e.g. CTCF site in mammals(3)), so far unidentified in yeast. Transcription activation can also interfere with apparition of cohesin dependent DNA structures that do not originate immediately at sites of transcription activated genes such as at the *GAL1-7,10* locus suggesting that transcription may interfere with cohesin mediated DNA loop during expansion process, by blocking the path to cis-translocating cohesin rather than powering DNA loop expanding cohesin. Not only cohesin was previously shown to compact transcriptionally inactive bacterial chromosomes *in vivo* (47), but also cohesin can extend DNA loops over long distances, encompassing multi-oriented genes, in the absence of Pds5 (19), indicating that polymerases are not required to either orient or push cohesin rings during DNA loop expansion process. To test this further, using a Pds5 mutant, we performed a kinetics showing how the induced *GAL7-10/1* loci temporarily halts the expansion of centromeric based DNA loops on the arm encompassing the GAL locus. The GAL block, at which *de novo* DNA loops emerge, are eventually gradually bypassed, showing that transcription represents a semi-permissive roadblock to cohesin mediated DNA loop expansion.

### A small proportion of cohesin may engage in DNA looping

Three pools of cohesin have been described so far, involved in i) DNA looping, ii) sister chromatids cohesion and iii) topological entrapping of a single chromatid (19,48–51). As cohesin ChIP-seq deposition profile does not discriminate between these populations, the relative proportions of cohesin involved in each pool, including the one involved in DNA looping, remains unknown. Our data shows that a small quantity of cohesin is blocked at the 3’ ends of active *GAL7-10/1* genes. This suggests that the overall amount of cohesin involved during the course of DNA looping may be relatively small. It is also possible that only few cohesin rings are required to hold DNA loops.

### How does transcription slow down the cohesin mediated DNA loop expansion process?

At least two models can explain how transcription positions DNA loops at the 3’ end of activated *GAL* genes. It is possible that the RNA polymerase dissociates the incoming DNA loop expanding cohesin from the gene body, and thus limits their position to the 3’ end of activated genes. RNA polymerase may also collide with the cohesin and then pushes it to the 3’end position of the gene. The latter scenario is the most likely, as evidenced by the effect of activating transcription of the *GAL2* locus in G2-arrested cells. Indeed, previous studies have revealed that RNA polymerase induces a shift of cohesin from the 5’-end to the 3’-end of activated *GAL2* even in the absence of *de novo* loading (22). In addition, we showed that cohesin displacement is accompanied by a decrease in the corresponding loop size. Taken together, these data show that RNA polymerase is a mobile barrier that limits the size of DNA loops.

We revealed DNA looping can be maintained at the 3’ end of oriented genes (*GAL1*) or divergent (*GAL7-10*) genes. Nevertheless, the base of most chromatin loops along budding yeast chromosomes are positioned between active convergent genes, suggesting that in addition to RNA polymerase, whose effect can be bypassed, other mechanism(s) independent of a specific DNA sequence, are required to maintain DNA loops at those specific loci. It is therefore likely that region in between convergent genes contain specific chromatin features or topology that favour the stabilization of the loops.

## Acknowledgements

This research was supported by the European Research Council under the Horizon 2020 Program (ERC grant agreement 771813) to RK, Agence Nationale pour la Recherche (ANR-22-CE12-0013-01) to FB and RK. FB also received support from the Fondation ARC pour la Recherche sur le Cancer (ARCPJA2022060005240) and the Comité de l’Occitanie de la Ligue Nationale contre le Cancer. NB was supported by the Ministère de l’Enseignement Supérieur and la Ligue Nationale contre le Cancer and Le comité départementale de la Ligue contre le Cancer de la Moselle. CC was supported by Pasteur-Roux-Cantarini fellowship.

We thank all our colleagues from the laboratories régulation spatiale des génomes and organization du noyau for helpful comments on the manuscript, K. Nasmyth’s laboratory and B. Albert for strains and K. Shirahige for the Smc3 antibody.

## Authors Contributions

Conceptualization: CC, NB, AC, RK and FB. Methodology: NB, CC, AC, OG, FB, RK. Investigation: NB, CC, FB. Formal analysis: CC, NB, AC. Data Curation: NB, CC, AC, FB, RK. Visualization: NB, CC, AC FB, RK. Writing: NB, CC, RK, FB. Supervision: FB, RK. Funding acquisition: FB, RK.

## Declaration of interest

The authors declare no competing interests.

## Figure Legends

**Supplementary Figure 1.**
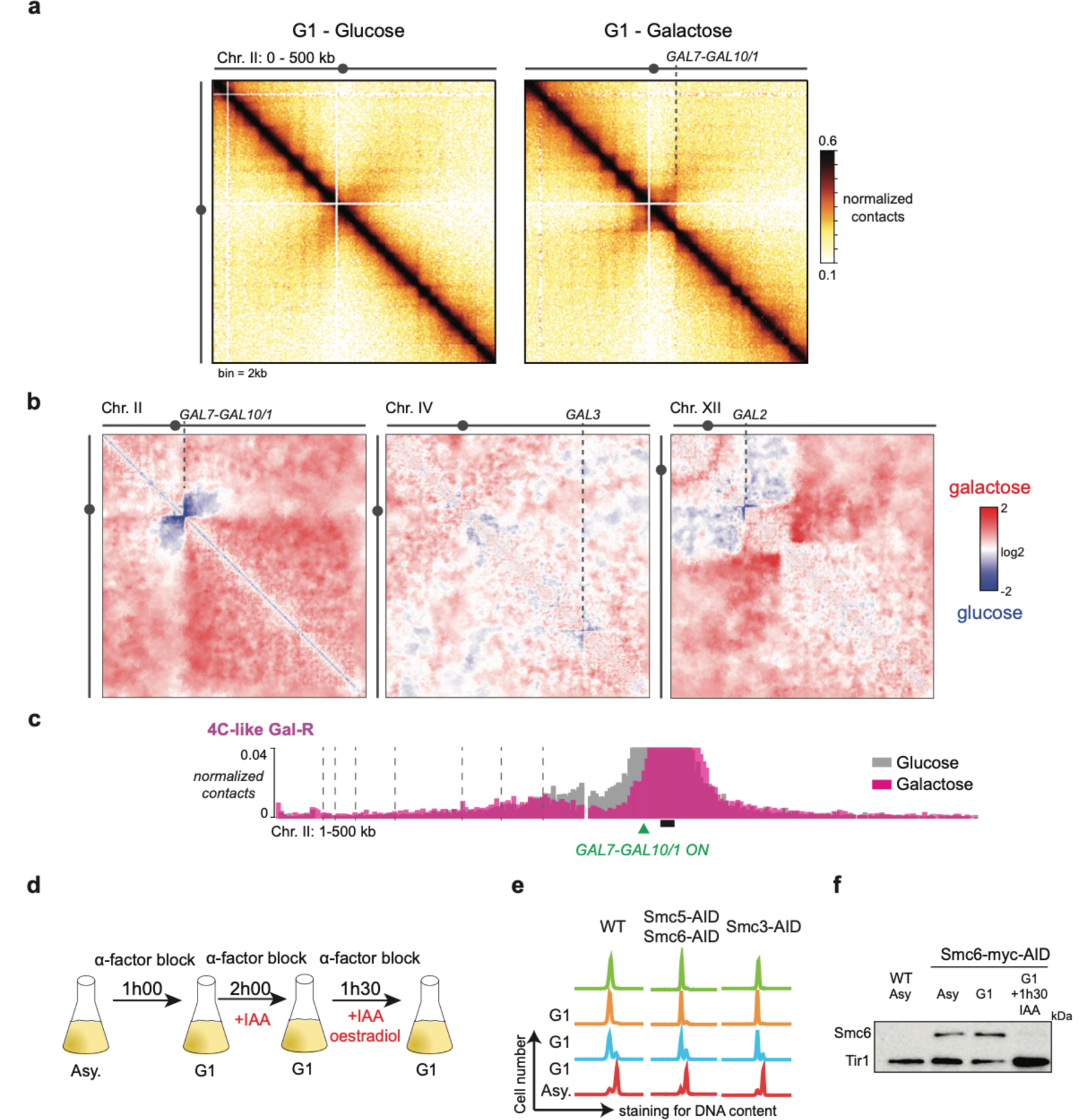
a) Hi-C contact maps (bin: 2kb) of a section of chromosome II (1-500kb) containing the GAL7-10/1 locus in cells synchronized in G1 in either glucose (left) or galactose (right). b) Ratio of contact maps obtained from G1 cells cultured in either glucose or galactose medium and processed through serpentine as described in the material and methods: chromosomes II (left), IV (middle) and XII (right). c) A virtual 4C plot with a viewpoint (anchor) just on the right side the GAL7-10/1 locus of cells in either glucose (grey) or galactose (pink). d) Schematic representation of the experimental protocol followed to induce GAL genes using oestradiol**AFTER** auxin-mediated SMC degradation (related to figure 1H). e) Synchronisation is controlled with by flow cytometry. Asy: asynchronous (related to figure 1H). f) Control of SMC degradation with a western blot (related to figure 1H).

**Supplementary Figure 2.**
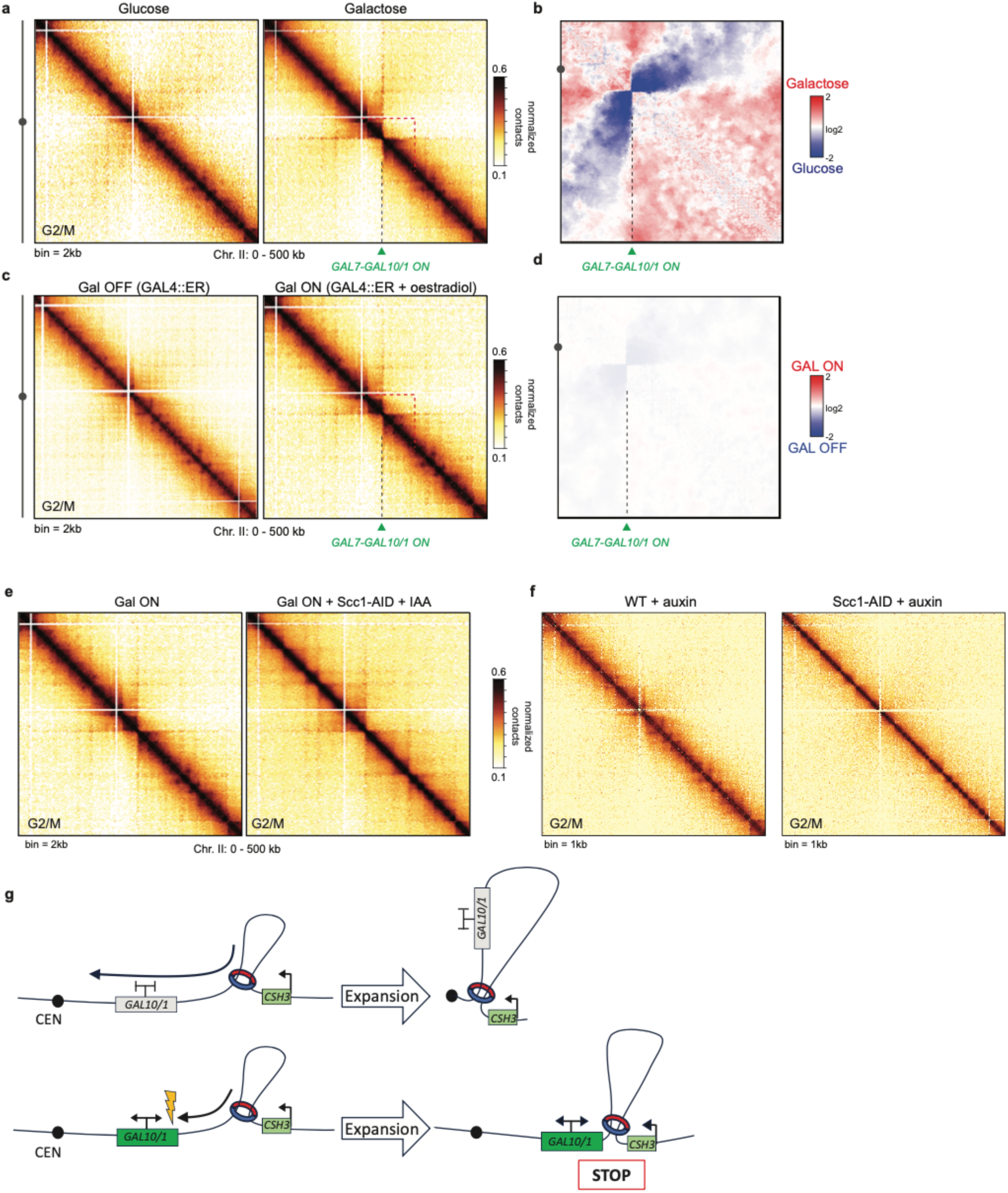
a) Hi-C contact maps (bin: 2kb) of a section of chromosome II containing the GAL7-10/1 locus in cells synchronized in G2 in either glucose or galactose containing medium. b) Ratio of the maps from panel **a** using serpentine as described in material and methods, showing chromosome II. c) Hi-C contact maps (bin: 2kb) demonstrating the effect of activation of the GAL7-GAL10/1 locus in G2 with oestradiol in cells. d) Ratio of the maps from panel **c** using serpentine as described in material and methods, showing chromosome II e) Hi-C contact maps (bin: 2kb) demonstrating the effect of activation of the GAL7-GAL10/1 locus in G2 with oestradiol in cells either lacking (GAL ON + Scc1-AID + auxin) cohesin. f) Hi-C contacts maps showing effect of Scc1 inactivation (Scc1-AID + auxin) on chromosome organization. g) Illustration of the mechanism by which transcription interferes on loop expansion.

**Supplementary Figure 3.**
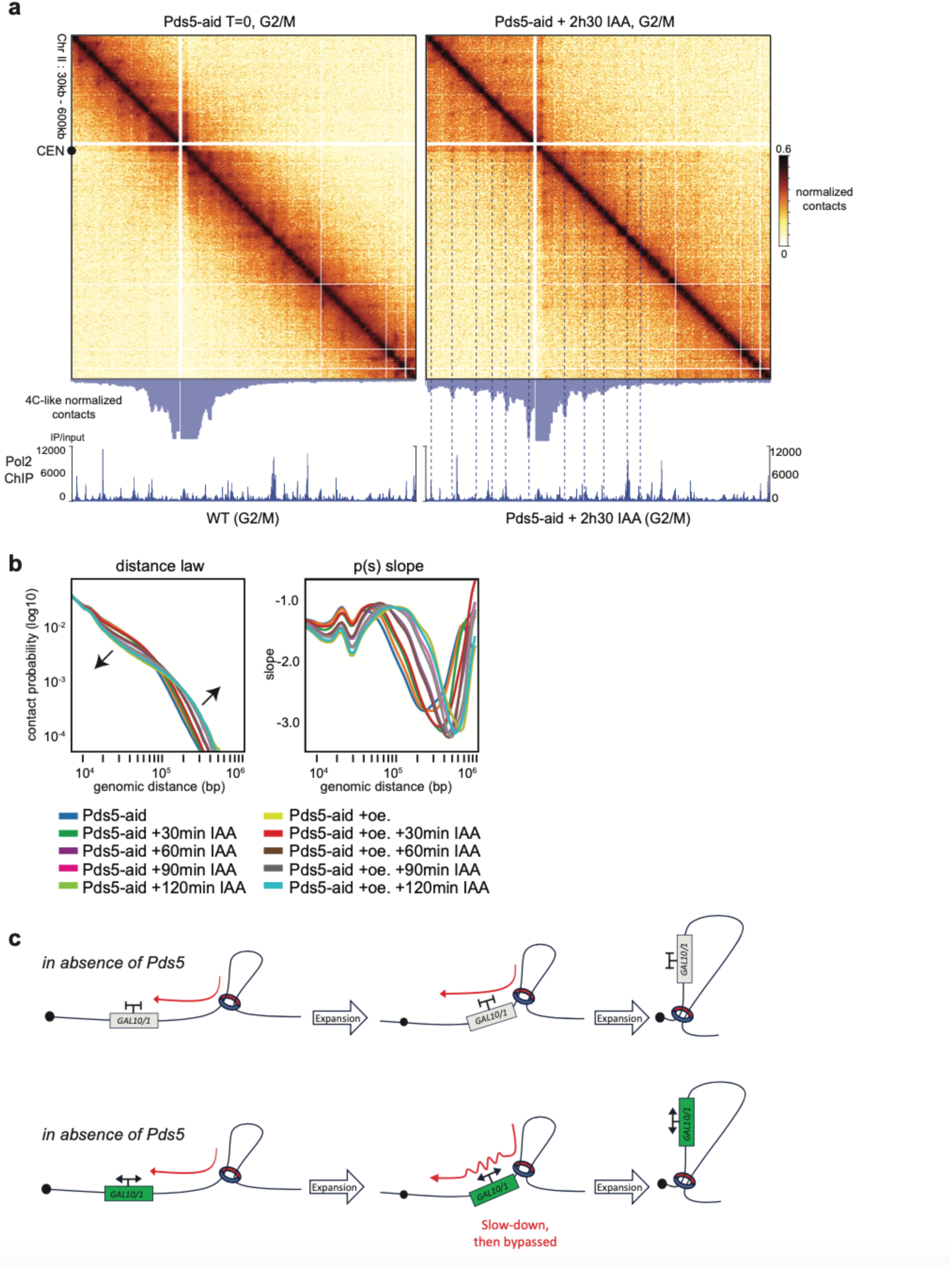
a) Hi-C contact maps (bin: 2kb), and corresponding RNA Polymerase II ChIP-seq profiles, of a section of chromosome II containing the *GAL7-10/1 locus* in in presence (left) or after depletion of Pds5 mediated by IAA (right). A virtual 4C plot with a viewpoint centred on the centromere is shown for each map. b) Contact frequency curves P(s), representing average contact frequency as a function of genomic separation (bp), and their respective derivative curves throughout Pds5 depletion from 0min to 120min after IAA addition. c) Model explaining how Gal activation slows-down the expansion of the long (CEN loop) induced by Pds5 depletion.

**Supplementary Figure 4.**
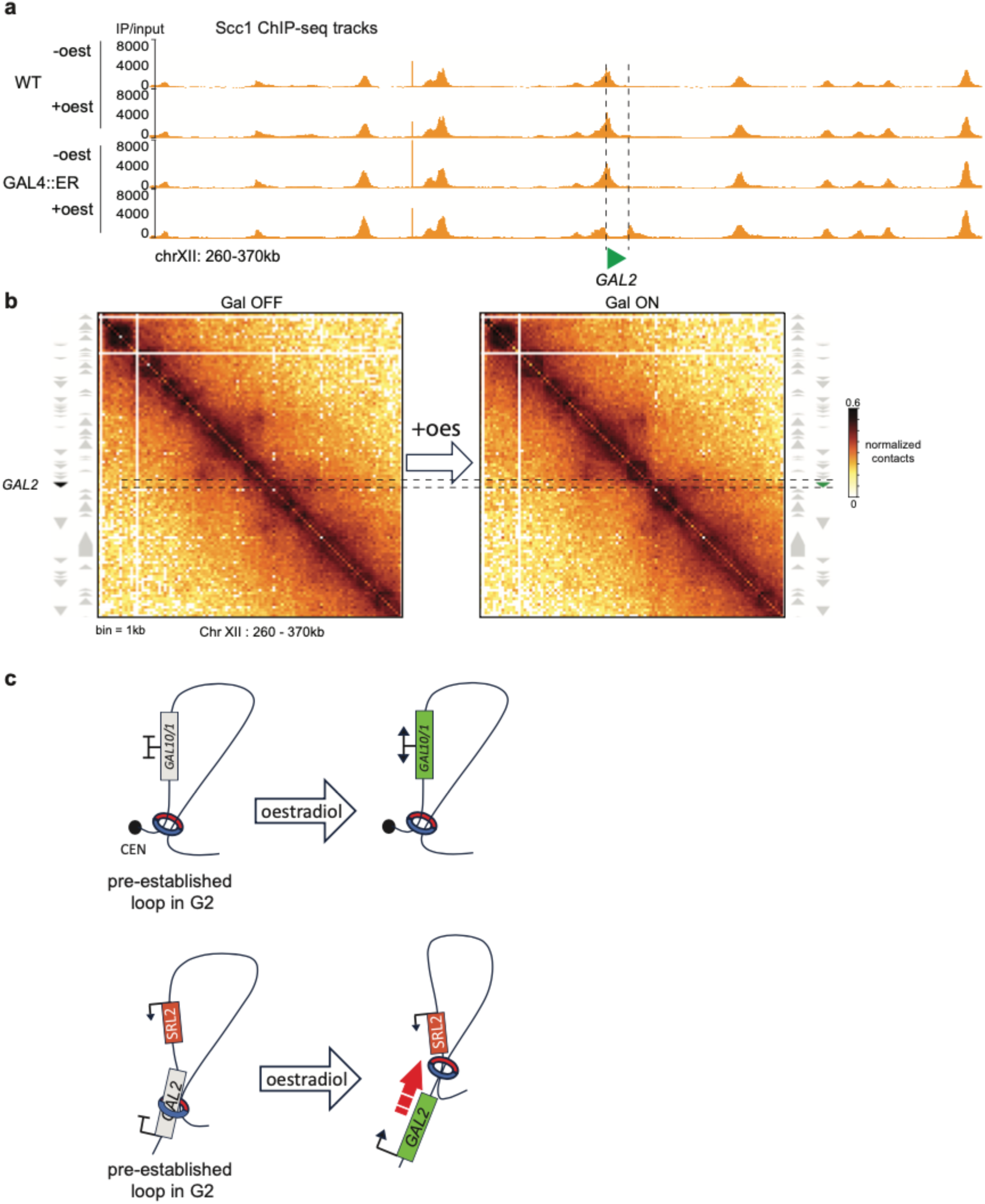
a) Scc1 ChIP-seq of a section of chromosome XII in cells arrested in G2 before (1st and 3rd tracks) and after adding oestradiol (2sd and 4th tracks). Dotted lines represent 5’ and 3’ of Gal2 gene. b) Contact maps (bin 1kb) of a section of chromosome XII in cells arrested in G2 in absence (left) or in presence (right) of GAL activation. Dotted lines represent 5’ and 3’ of Gal2 gene. c) Illustration explaining effect of transcription on the maintenance of DNA loop in G2/M.

**Table 1.** Yeast strain used in this study.

| Strain name | Background | genotype | Figure | Ref |
| --- | --- | --- | --- | --- |
| W303-1A | W303 | MATa | Fig 1, Fig 2, Fig 4, sup Fig 1, sup Fig 2, Sup Fig 3 | Ralser, M. et al. The *Saccharomyces cerevisiae* W303-K6001 cross-platform genome sequence: insights into ancestry and physiology of a laboratory mutt. *Open Biol.* 2, 120093 (2012). |
| BEN15 | W303 | MATa ura3:GAL4DBD-ER-Msn2-AD:URA3 | Fig 4 | Pincus et al. 2014 |
| FB220-1b | W303 | MATa ura3:GAL4DBD-ER-Msn2-AD:URA3, his3::ADH1promoter-OsTIR1-9myc::HIS3 | Fig 2, sup fig 2 | This study |
| FB219-2a | W303 | MATa ura3:GAL4DBD-ER-Msn2-AD:URA3, Scc1-PK3-aid::KanMX4, his3::ADH1promoter-OsTIR1-9myc::HIS3 | sup fig 2, sup Fig 4 | This study |
| FB220-2a | W303 | MATa his3::ADH1promoter-OsTIR1-9myc::HIS3, Pds5-AID::KanMx | Fig 3, sup Fig 3 |  |
| FB220-8c | W303 | MATa ura3:GAL4DBD-ER-Msn2-AD:URA3, his3::ADH1promoter-OsTIR1-9myc::HIS3, Pds5-AID::KanMx | Fig 3, sup Fig 3 | Dauban et al. 2020 |
| FB217-13C | W303 | MATa his3::ADH1promoter-OsTIR1-9myc::HIS3, SCC1-HA6::HIS3, ura3:GAL4DBD-ER-Msn2-AD:URA3, Pds5-AID::KanMx | Fig 3 | This study |
| FB222-1c | W303 | MATa his3::ADH1promoter-OsTIR1-9myc::HIS3, SCC1-HA6::HIS3, Pds5-AID::KanMx | Fig 3 | This study |
| FB218-4a | W303 | MATa SCC1-HA6::HIS3 | Fig 2, Fig 4, sup Fig 4 | This study |
| FB218-8D | W303 | MATa SCC1-HA6::HIS3, ura3:GAL4DBD-ER-Msn2-AD:URA3 | Fig 2, Fig 4, sup Fig 4 | This study |
| KN25532 | Candida Glabrata | Scc1-HA3::NatMX | Fig 2 | Petela et al Mol Cell |
| KN23308 | Candida Glabrata | Scc1-pk9::NatMX | Fig 1 | Hu et al, NAR |
| yNB30.2-14b | W303 | MATa SCC1-PK9::KanMX | Fig 1 | This study |

**Table 2.**
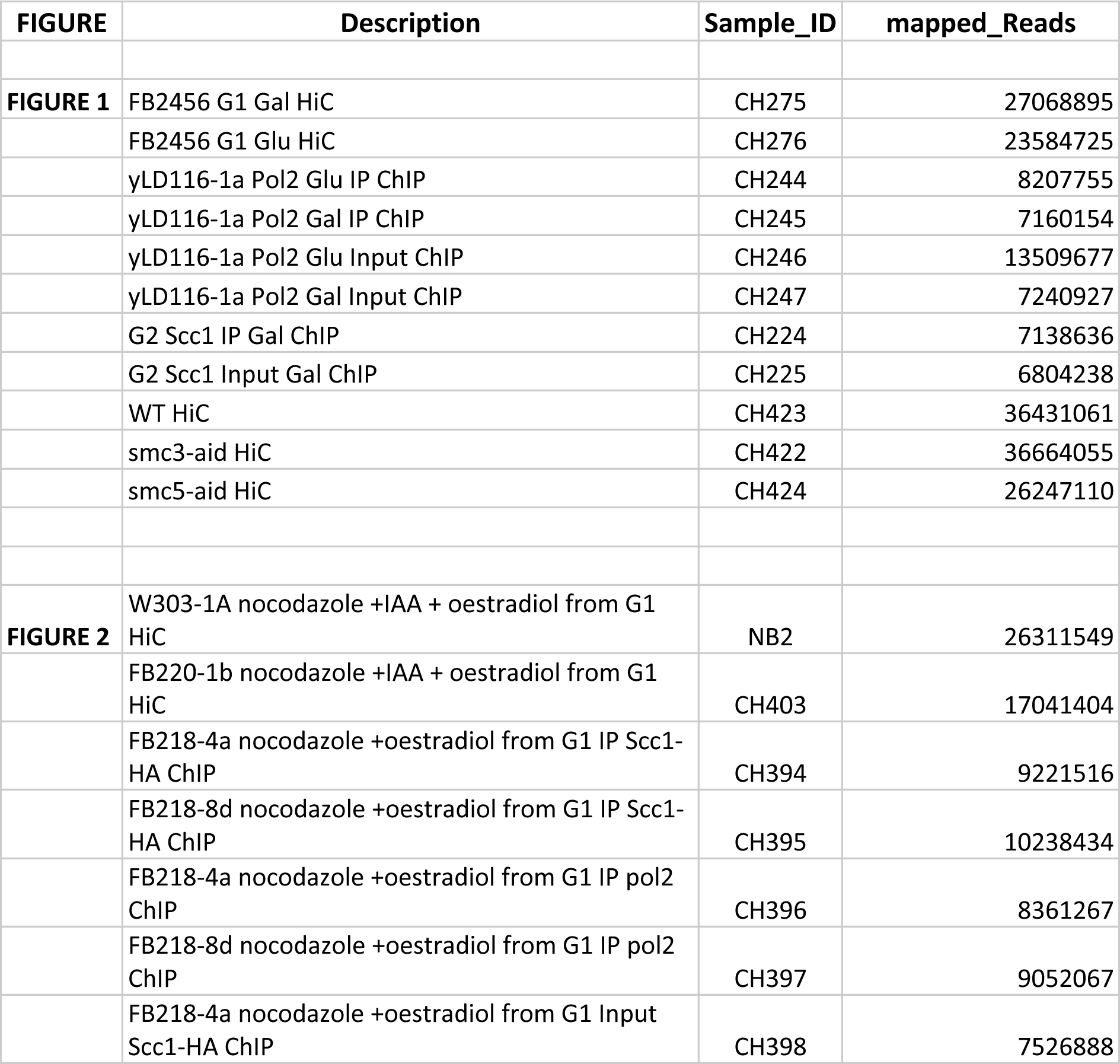
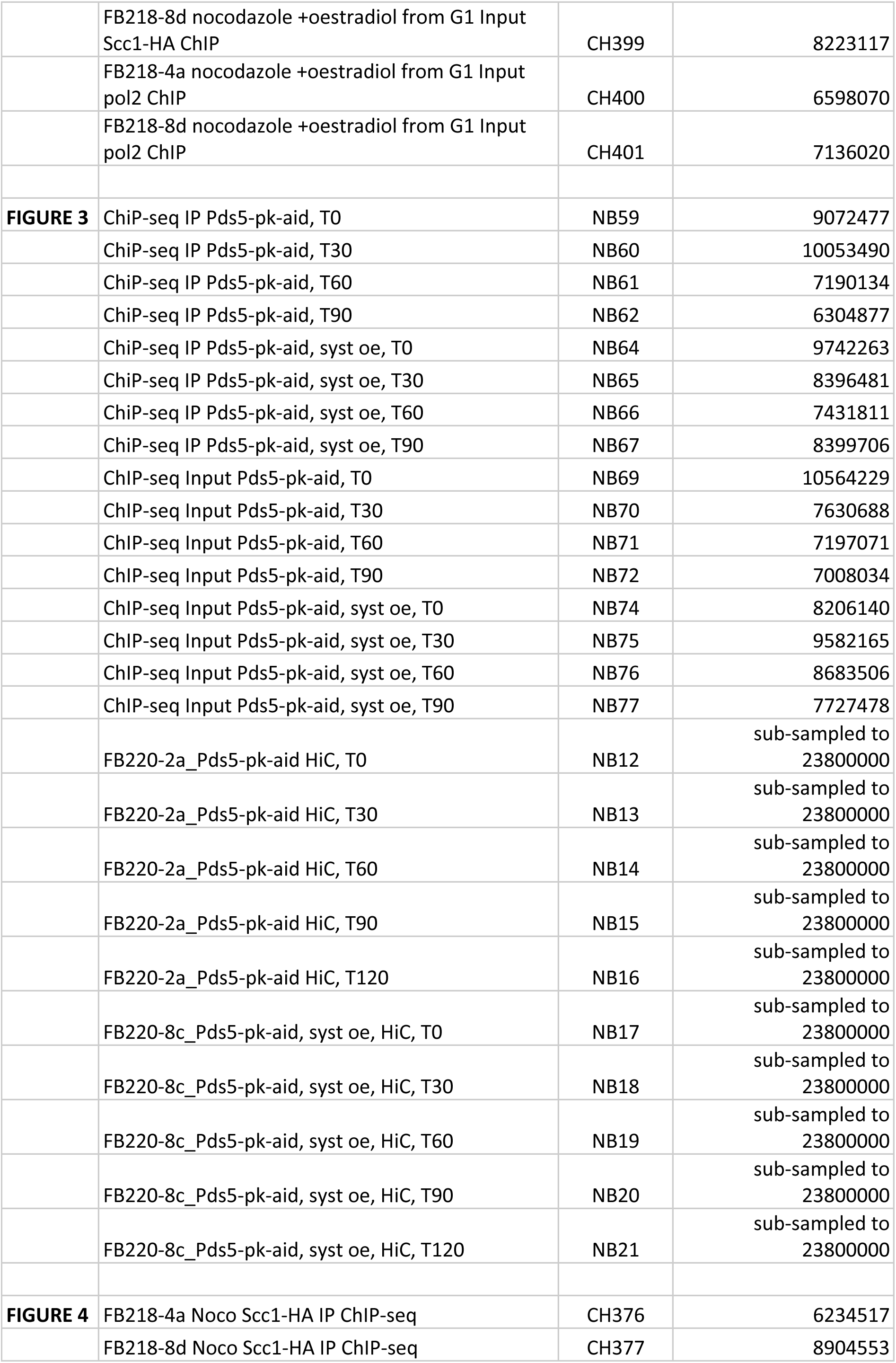
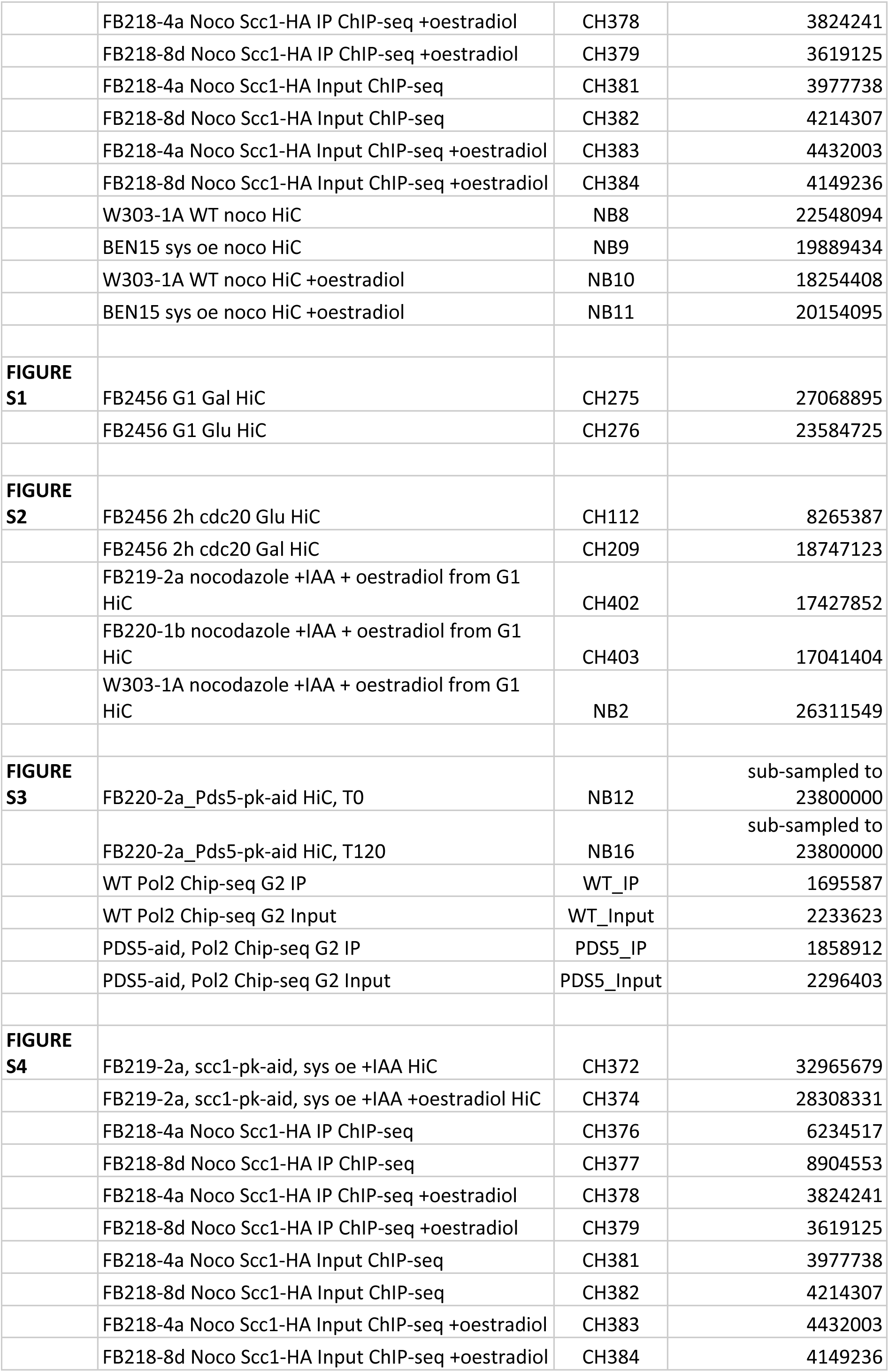
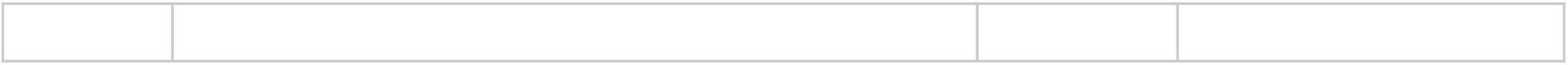
Table of reads.

## Material and Methods

### Data Availability

Accession number for processed data for all figures: GSE232323.

Raw sequences for all figures are accessible on SRA database through the following accession number: PRJNA971611

### Reference genomes

a. *S. cerevisiae* W303 (52); *C. glabrata* (53): http://www.candidagenome.org/download/sequence/C_glabrata_CBS138/current/

Strains used in this study are available upon request.

### Code Availability

Programs involved in the study are listed below.

HiCstuff (www.github.com/koszullab/hicstuff) version 3.0.1 (34)

Serpentine (www.github.com/koszullab/serpentine; version 0.1.3 (34)

Bowtie2 (version 2.3.4.1 available online at http://bowtie-bio.sourceforge.net/bowtie2/)(54) Samtools (version 1.9 available online at http://www.htslib.org/download/)(55)

Bedtools (version 2.29.1 available online at https://bedtools.readthedocs.io/en/latest/content/installation.html)(56)

Cooler (version 0.8.7-0.8.11 available online at https://cooler.readthedocs.io/en/latest/)(57) bamCoverage (version 3.4.1) https://deeptools.readthedocs.io/en/develop/content/tools/bamCoverage.html)(58)

### Experimental procedures

#### Media culture conditions and synchronisation

All strains used in this study are derivative of W303 and are listed in the Table “Strain list”. All strains were grown overnight at 30°C or 25°C in 150mL of suitable media supplemented with either 2% glucose or 2% galactose where indicated to attain 4,2x10^8 cells. Yeast cells were synchronised in G1 by adding of α-factor (Antibodies-online, ABIN399114 or Proteogenix, WY-13) in the media every 30 min during 2h30 (1µg/mL final).

To arrest cells in metaphase, yeast cells were synchronised in G1 by adding of α-factor in the media every 30 min during 2h30 (1µg/mL final). After G1 arrest, cells were washed twice in fresh YP and released in rich medium (YP): 1%bacto peptone (Difco), 1%bacto yeast extract (Difco), supplemented with either 2% glucose or 2% galactose where indicated and containing Nocodazole (Sigma-Aldrich, M1404-10MG). Auxin (2mM final) (Sigma-Aldrich, I3750) and/or oestradiol (100nM final) (Sigma-Aldrich, E2758) was added to the media when indicated.

#### Flow cytometry

To verify cell cycle arrest and synchronisation, 1mL of cells culture were fixed in ethanol 70% and stored overnight at −20°C. Pellet was incubated with 50mM Tris-HCl (pH 7,5) and 5µL RNase A (10mg/mL) overnight at 37°C. Cells were pelleted and re-suspended in 400µL of FACS buffer (1mg/mL propidium iodide (Fisher, P3566), Tris-HCl, NaCl, MgCl2) and incubated at 4°C. Cells were sonicated with 60% output for 10 seconds.

Flow cytometry was performed on a CytoFLEX S (Beckman Coulter) and data were analysed using CytExpert 2.4 software, measuring 1000 events at a 30µL/min flow rate.

#### Protein extractions and acetylation assays

A pellet from 10mL of culture was frozen in liquid nitrogen and stored overnight at −20°C. Cell pellets were re-suspended in 100µL H2O and 20µL trichloroacetic acid (Sigma-Aldrich, T8657) (TCA). Then cells were broken by glass beads at 4°C and precipitated proteins were re-suspended in Laemmli buffer with 100mM DTT and Tris-HCl (pH 9,5). Proteins were extracted by cycles of 5 min heating at 80°C and 5min vortexing at 4°C. After centrifugation, extracted proteins were collected and froze at −20°C.

Eluates were analysed by SDS-page followed by western blotting with antibodies:

- Mouse anti Smc3-K113Ac (Beckouet et al., 2010) (ab from K. Shirahige laboratory, clone H2) used at dilution 1:1000 for Western Blot
- Mouse anti-V5 tag (Bio-Rad, MCA1360) used at dilution 1:5000 for Western Blot
- Mouse anti-HA (Abcam, ab1424, 12CA5), used at dilution 1:1,000 for Western Blot
- Mouse anti-pgk1 (22C5D8) (Invitrogen, 459250) used at dilution 1:5000 for Western Blot
- Anti-mouse IgG, HRP conjugate (Promega, W4021) used at dilution 1:5000 for Western Blot

Blots were revealed using ChemiDoc Touch Imaging System: Image Lab 6.0.

The ratio (relative quantity of acSmc3/Pgk1) was calculated using the Volume tool of Image Lab: Background subtraction method: Local, Quantity regression method: Point to Point through origin.

#### Calibrated ChIP-sequencing

Cells were grown exponentially to OD600 = 0.5. In triplicates, 15 OD600 units of *S. cerevisiae* cells were mixed with 3 OD600 units of *C. glabrata* to a total volume of 45 mL and fixed with 4mL of fixative solution (50 mM Tris-HCl, pH 8.0; 100 mM NaCl; 0.5 mM EGTA; 1 mM EDTA; 30% (v/v) formaldehyde) for 30 min at room temperature (RT) with rotation. The fixative was quenched with 2mL of 2.5M glycine (RT, 5 min with rotation). The cells were then harvested by centrifugation at 3,500 rpm for 3 min and washed with ice-cold PBS. The cells were then resuspended in 300 mL of ChIP lysis buffer (50 mM HEPES KOH, pH 8.0; 140 mM NaCl; 1 mM EDTA; 1% (v/v) Triton X-100; 0.1% (w/v) sodium deoxycholate; 1 mM PMSF; 2X Complete protease inhibitor cocktail (Roche)) and transfer in tubes 2mL containing glass beads before mechanical cells lysis. The soluble fraction was isolated by centrifugation at 2,000 rpm for 3min then transferred to sonication tubes and samples were sonicated to produce sheared chromatin with a size range of 200-1,000bp. After sonication the samples were centrifuged at 13,200 rpm at 4°C for 20min and the supernatant was transferred into 700µL of ChIP lysis buffer. 80µL (27µl of each sample) of the supernatant was removed (termed ‘whole cell extract [WCE] sample’) and store at −80°C. 5ug of antibody (anti-HA or anti-Pol2 8WG16) was added to the remaining supernatant which is then incubated overnight at 4°C (wheel cold room). 50µL of protein G Dynabeads was then added and incubated at 4°C for 2h. Beads were washed 2 times with ChIP lysis buffer, 3 times with high salt ChIP lysis buffer (50mM HEPES-KOH, pH 8.0; 500 mM NaCl; 1 mM EDTA; 1% (v/v) Triton X-100; 0.1% (w/v) sodium deoxycholate;1 mM PMSF), 2 times with ChIP wash buffer (10 mM Tris-HCl, pH 8.0; 0.25MLiCl; 0.5% NP-40; 0.5% sodium deoxycholate; 1mM EDTA;1 mMPMSF) and 1 time with TE pH7.5. The immunoprecipitated chromatin was then eluted by incubation in 120µL TES buffer (50mM Tris-HCl, pH 8.0; 10 mM EDTA; 1% SDS) for 15min at 65°C and the supernatant is collected termed ‘IP sample’. The WCE samples were mixed with 40µL of TES3 buffer (50mM Tris-HCl, pH 8.0; 10mM EDTA; 3% SDS). ALL (IP and WCE) samples were de-cross-linked by incubation at 65°C overnight. RNA was degraded by incubation with 2µL RNase A (10 mg/mL) for 1h at 37°C. Proteins were removed by incubation with 10µL of proteinase K (18 mg/mL) for 2h at 65°C. DNA was purified by a phenol/Chloroform extraction. The triplicate IP samples were mixed in 1 tube and libraries for IP and WCE samples were prepared using Invitrogen TM Collibri TM PS DNA Library Prep Kit for Illumina and following manufacturer instructions. Paired-end sequencing on an Illumina NextSeq500 (2x35 bp) was performed. For analysis, Bowtie2 was used for two rounds of alignments, first on *C. glabrata* (CBS138) and then on *S. cerevisiae* allowing the generation of an alignment of IP and WCE that exclusively mapped on *S. cerevisiae* (an vice-versa for *C. glabrata*). The obtained SAM file was converted into a BAM file, sorted and indexed using Samtools. ChIP-seq profiles were then normalised by the number of million sequences and converted into BigWig using bamCoverage. Profiles were multiplied by the ORi factor (WCEglabrataIPcerevisiae / WCEcerevisiaeIPglabrata, in which WCE glabrata and IPglabrata correspond to the number of paired reads that mapped uniquely on *C. glabrata* genome and same for *S. cerevisiae* reads) using Integrated Genome Browser.

#### Hi-C procedure and sequencing

Cell fixation with 3% formaldehyde (Sigma-Aldrich, Cat. F8775) was performed as described in Dauban et al. 2020 and Piazza et al. 2020 (59)(19). Quenching of formaldehyde with 300 mM glycine was performed at 4°C for 20 min. Hi-C experiments were performed with a Hi-C kit (Arima Genomics) with a double DpnII + HinfI restriction digestion following manufacturer instructions. Briefly, samples were permeabilised by sequentially adding lysis buffer (15min at 4C), conditioning solution (10min at 62C) and stop solution 2 (15min at 37C) to the samples. DNA was digested using a mix of buffer A, DpnII, and Hinf1 (45min at 37C followed by 20min at 65C). DNA was repaired and biotin was added by adding a mix of buffer B and enzyme B for 45min at room temperature. DNA was re-ligated by adding a mix of buffer C and enzyme C during 15min at room temperature. Samples were then digested by protease and de-crosslinked by adding a mix of buffer D, enzyme D and buffer E during 30 min à 55°C followed by 90 min at 68°C. Samples were purified using AMPure XP beads (Beckman A63882), recovered in 120ul H2O and sonicated using Covaris (DNA 300bp). Preparation of the samples for paired-end sequencing on an Illumina NextSeq500 (2x35 bp) was performed using Invitrogen TM Collibri TM PS DNA Library Prep Kit for Illumina and following manufacturer instructions.

Processing of the reads, computation of contact matrices, and generation of contact maps Reads were aligned and the contact data processed using Hicstuff, available on Github (https://github.com/koszullab/hicstuff). Briefly, pairs of reads were aligned iteratively and independently using Bowtie2 (54) in its most sensitive mode against the *S. cerevisiae* W303 reference genome. Each uniquely mapped read was assigned to a restriction fragment. Quantification of pairwise contacts between restriction fragments was performed with default parameters: uncuts, loops and circularization events were filtered as described previously (40).

PCR duplicates (defined as paired reads mapping at exactly the same position) were discarded. Contact maps from independent replicates were generated with the “view” function of Hicstuff, merged and normalized using the merge and balance functions of Cooler. Bins were set at 1, exp0.2 transformed, and rendered.

#### Computation of the contact probability as a function of genomic distance

Computation of the contact probability as a function of genomic distance P(s) and its derivative have been determined using the “distance law” function of Hicstuff with default parameters, averaging the contact data of entire chromosome arms. P(s) from two independent replicates were determined and plotted.

#### Generation of the ratio maps using serpentine

Ratio maps were generated with Serpentine (34). Briefly, comparison between pairs of 1-kb contact maps was performed using the default threshold parameters and the detrending constant was determined automatically by Serpentine for each comparison.

#### Virtual 4C profiles

4C-like profiles were generated on 2kb binned contact maps for a region of interest using an 8kb window as bait.

#### Loop spectrum using chromosight “quantify” mode

The Loop Spectrums were computed as explained in (34). First, peaks of RNA PolII were extracted from Rpb3 ChIP-seq profiles generated during in log phase (60) with homemade python codes. Enrichment in contacts (i.e. “loop scores”) between all pairs of loci enriched for a peak in Rpb3 were computed using Chromosight, and the average loop scores were computed for different sizes of loops on Micro-C contact data from (60), (SRR13736654). A locally weighted scatterplot smoothing (lowess) was then applied using scikit-misc package. The pileups plots of either windows centred on RNA PolII peaks along the diagonal, or of pairs of windows centred on distant RNA PolII peaks (60 kb to 460 kb), were computed for five different groups of RNA peaks sorted by the transcription level (with ChIP-seq of Rpb3 data of (60)). The different groups of transcription levels were determined by computing the distribution of ChIP-Seq intensities for all 2 kb bins of the *S. cerevisiae* genome. Then the five groups were defined with the following limits: [0:0,5],[0,5:1,0],[1:1,25],[1,25:1,5] and [1,5:15] corresponding to regions from low to high RNA PolII levels, respectively.

Pileup plots during the mitotic cell cycle were generated with 2 different group of pairs: pairs of cohesin peaks between 10 kb and 50 kb and pairs of RNA Pol II peaks between 60 kb and 460 kb with the contact data from (39), (SRR11893107). Pileup plots for SMC mutants were generated with the same group of pairs of cohesin peaks between 10 kb and 50 kb.

## Notes

### Competing Interest Statement

The authors have declared no competing interest.

